# Conservation of Nuclear Receptor DAF-12 with Evidence for Ligand Diversification in Plant-Parasitic Nematodes

**DOI:** 10.64898/2026.09.03.748874

**Authors:** Yanjie Liu, Chih-Wei Fan, Zhu Wang, George Wendt, Lauren G. Zacharias, Jeffrey G. McDonald, Yunhai Li, Shi-Yi Shen, H. Eric Xu, James J. Collins, Steven Kliewer, Melissa G. Mitchum, David J. Mangelsdorf, Thomas P. Mathews

## Abstract

Plant-parasitic nematodes cause extensive crop losses that threaten global food security and create substantial economic hardship, yet strategies for their control remain limited. Although plant-parasitic nematodes have life cycles that mirror those of their free-living and animal-parasitic relatives, comparatively little is known about the signals that drive life-cycle progression. Dafachronic acids (DAs) are steroid hormones that control life cycle by activating DAF-12 nuclear receptors in *Caenorhabditis elegans* and animal-parasitic nematodes. To define whether an analogous pathway exists in plant-parasitic nematodes, we investigated the responsiveness of DAF-12 orthologs from six plant-parasitic nematode species to Δ7-DA, an endogenous DAF-12 ligand in *C. elegans* and several animal-parasitic nematodes. We find that whereas some plant-parasitic nematode receptors are activated by Δ7-DA, others are not.

Structural modeling of all six ligand-binding domains, together with docking of Δ7-DA into each model, indicates that responsiveness is governed by a combinatorial ligand-recognition network surrounding the ligand carboxylate rather than by any single contact residue. We further demonstrate that egg and second-stage juvenile extracts from the soybean cyst nematode, *Heterodera glycines*, lack detectable DA but contain distinct lipophilic fractions that activate its DAF-12 ortholog. Together, these findings suggest that many plant-parasitic nematodes utilize endogenous DAF-12 ligands that are structurally distinct from DA. Uncovering modulators of this pathway could lead to new strategies to combat nematode-driven crop loss.

## Introduction

Plant-parasitic nematodes are among the most destructive agricultural pathogens, threatening food security and causing billions of dollars in crop losses worldwide each year (1). Despite their enormous impact, relatively little is known about the pathways that govern their development and life-cycle progression. In the free-living nematode *C. elegans*, developmental outcomes are orchestrated by the nuclear receptor DAF-12, which integrates environmental and metabolic cues to regulate reproductive development (2). Orthologs of DAF-12 are conserved throughout the phylum Nematoda, where they control entry into and exit from infective larval stages in animal-parasitic nematodes. These findings have established DAF-12 as a master developmental regulator and an attractive target for controlling parasitic nematodes (3).

Bile acid-like steroid hormones known as dafachronic acids (DAs) function as endogenous DAF-12 ligands in *C. elegans* and several animal-parasitic nematodes (4–8). In *C. elegans*, Δ4-DA and Δ7-DA were initially identified as endogenous ligands (7), and subsequent comparative metabolomics identified Δ1,7-DA as a major endogenous ligand (6). Δ4-DA and Δ7-DA have also been identified in animal-parasitic nematodes, with their relative abundance varying among species (4; 5; 8). Crystal structures of the DAF-12 ligand-binding domain (LBD) from the human parasite *S. stercoralis* bound to Δ7-DA defined the molecular basis of this recognition: the C27 carboxylate of the ligand is coordinated by a network centered on a conserved arginine (R599), the C3-keto group is read out by a glutamine (Q637) on helix 8, and ligand binding stabilizes the C-terminal AF-2 helix that recruits transcriptional coactivators (9). The identification of DAF-12 ligands in *C. elegans* and animal-parasitic nematodes raised the intriguing possibility that plant-parasitic nematodes employ a comparable hormonal mechanism to regulate their life cycles.

Much of our current understanding of DAF-12 biology and DA signaling stems from the pioneering work of Dr. David Mangelsdorf and colleagues. Over the past two decades, David’s laboratory discovered DAs as endogenous DAF-12 ligands, elucidated the structural basis for ligand recognition, demonstrated the conservation of DAF-12 signaling across numerous parasitic nematode species, and established the pathway as a promising therapeutic target for controlling nematode infections (7–11). These studies transformed DAF-12 from an orphan nuclear receptor into one of the best understood regulators in nematode biology and laid the foundation for exploring developmental signaling in parasitic species. It is therefore fitting that this study, which extends these questions to plant-parasitic nematodes, is dedicated to David in recognition of his profound and lasting contributions to the field.

Here, we examine the responsiveness of DAF-12 orthologs from representative plant-parasitic nematodes to Δ7-DA and investigate endogenous DAF-12 ligands in the soybean cyst nematode, *Heterodera glycines*. We find that some plant-parasitic DAF-12 receptors are activated by Δ7-DA whereas others are not and have corresponding structural differences in their LBDs. We further identify lipophilic fractions from *H. glycines* egg and second-stage juvenile (J2) extracts that activate DAF-12 despite the absence of detectable DA. Together, these findings demonstrate conservation of DAF-12 in plant-parasitic nematodes while providing evidence for diversification of its endogenous ligands.

## Materials and Methods

### Cell-based reporter assays for DAF-12 transcriptional activity

DAF-12 transcriptional activity was measured using glucocorticoid receptor (GR)/DAF-12 chimeric receptors expressed from a CMV-driven mammalian expression vector (pCMX). Chimeras consisted of the N-terminal region and DBD of the human GR (amino acids 1-552; NCBI X03225), the hinge region of *C. elegans* DAF-12 (amino acids 184-487; F11A1.3a.1), the hinge and LBD of DAF-12 from the indicated species, and the C-terminal 10 amino acids of human SRC-1a (amino acids 1432-1441; RefSeq NM_003743.3). DNA fragments encoding predicted DAF-12 sequences were synthesized for *C. elegans* (F11A1.3a.1; RefSeq NP_001379300.1), *H. glycines* (Hetgly12168), *G. rostochiensis* (Gr22_v10_g7179_t1), *B. xylophilus* (BXY_1418300), *A. besseyi* (M3Y94_00792600), *Meloidogyne incognita* (Minc3s00350g10770), and *D. destructor* (Dd_05346). The hinge-LBD amino acid sequences incorporated into each chimera are provided in Supplementary Figure S1.

For transfection analyses, COS-7 cells were maintained in Dulbecco’s modified Eagle’s medium (DMEM; Thermo Fisher Scientific) supplemented with 10% (v/v) fetal bovine serum and 100 U/mL penicillin–streptomycin. Cells were seeded in 6-well plates and co-transfected with 200 ng pCMX-GR-DAF-12 expression plasmid, 600 ng MTV-firefly luciferase reporter plasmid (12), and 200 ng pNL1.1.PGK NanoLuc control plasmid (Promega) using X-tremeGENE HP DNA Transfection Reagent (Roche). Twenty-four hours after transfection, cells were replated into 384-well plates and treated for 17–24 h with vehicle, Δ7-dafachronic acid (Δ7-DA) or HPLC fractions (2%, v/v). Firefly and NanoLuc luciferase activities were measured using the Nano-Glo Dual-Luciferase Reporter Assay System (Promega), and reporter activity was expressed as firefly luciferase normalized to NanoLuc and then normalized to vehicle-treated controls to calculate fold activation. Dose-response curves were analyzed by nonlinear regression using three-parameter logistic models.

### Structural modeling and ligand docking

Structural models of the DAF-12 LBDs from *H. glycines*, *G. rostochiensis*, *B. xylophilus*, *A. besseyi*, *M. incognita* and *D. destructor* were generated with AlphaFold3 using the LBD sequences employed in the chimeric reporter constructs, so that the models correspond exactly to the proteins tested functionally. The 2.4 Å co-crystal structure of the *S. stercoralis* DAF-12 LBD bound to Δ7-dafachronic acid and an SRC-1 coactivator peptide (PDB 3GYU) was used as the reference for the active-state conformation (9). Predicted models were superposed on the reference structure over the LBD core, and Δ7-DA was redocked into each model using the ligand pose of 3GYU to define the search volume; resulting poses were energy-minimized and inspected for steric complementarity within the pocket. Electrostatic surfaces of the ligand-binding cavity were calculated and are displayed on a scale of −10 to +10 kT/e. Structure-guided sequence alignment of the seven LBDs was performed using the secondary-structure assignment of the *S. stercoralis* structure, and binding-site positions were assigned by correspondence to the crystallographically defined Δ7-DA contacts in 3GYU (T562, R599, T613 and Q637). Molecular graphics were prepared with ChimeraX and the sequence alignment figure was rendered with ESPript 3.0. Model coordinates are available from the authors on request.

### Phylogenetic and phylogenomic analyses

Sequence data and domain alignments: Protein sequences of 144 nematode DAF-12 homologs from 120 species (Table S1) were retrieved from WormBase ParaSite release WBPS19 (WS291) (13), using *C. elegans* DAF-12 (F11A1.3) as the reference. Three additional DAF-12 sequences (TTRE6457, Gr19_v10_g11452, ES5_v2.g15056) were identified manually by reciprocal BLAST (Table S2, green). The DAF-12 LBD was isolated and aligned from these sequences using MAFFT-LAST (14; 15) (default settings except minimum coverage set to 0.3) with the LBD from *C. elegans* DAF-12 (Table S3) as the reference sequence. The final LBD alignment comprised 135 sequences from 115 species (Table S4). Each species was assigned a single trophic mode — free-living, plant-parasitic, animal-parasitic with a vertebrate host, or animal-parasitic with an insect host — from published life-history descriptions. Taxonomic ranks follow the NCBI Taxonomy hierarchy and clade nomenclature follows the molecular framework of Blaxter and colleagues (16).

For maximum-likelihood phylogenetic inference, substitution models were selected and trees inferred with IQ-TREE 2 v2.4.0 (17). ModelFinder (18) selected Q.insect+G4 for the LBD by Bayesian information criterion (BIC = 35007.98, 256 free parameters), with four discrete gamma rate categories and shape parameter alpha = 1.03. The maximum-likelihood tree had a log-likelihood of -16790.27 (s.e. 818.24) and a total length of 28.39 substitutions per site, 49.6% of which lay on internal branches. Trees were rooted on *Plectus sambesii* (Plectida), the most distant outgroup available in the sampled set. Tree manipulation, rooting and clade-membership queries used ETE 3 (19). Branch support was estimated within the same run by two independent methods: 1000 ultrafast bootstrap replicates (UFBoot2) (20) with NNI optimization on bootstrap alignments, and the SH-aLRT test with 1000 replicates (21). Support is reported throughout as SH-aLRT / UFBoot. Following the recommendation of the UFBoot2 authors, a branch was considered strongly supported only when SH-aLRT >= 80 and UFBoot >= 95; branches with UFBoot 70-94 are described as moderately supported and those with UFBoot < 70 as unsupported. Tree results in a Newick format are available as File S1.

For the comparative genomic survey of DAF-36, whole predicted proteomes were obtained from WormBase ParaSite WBPS19 (13) for all 115 species represented in the LBD alignment, together with four additional species present in the proteome collection but not in the alignment, giving 120 proteomes. Each proteome was initially matched to the specific BioProject assembly from which the corresponding aligned DAF-12 sequence derived, so that gene-content calls and phylogenetic sampling refer to the same assembly rather than to different assemblies of the same species. To ensure DAF-36 survey completeness, we also searched all available assemblies of any species whose initial call was absent or unresolved: 11 of these species scored absent and had multiple assemblies, so all 27 of their assemblies were searched. Additional survey of NCBI identified another Clade 12 nematode (*Aphelenchus avenae*) that has a DAF-36 homolog, so it was also included in the genomic survey pipeline. A DAF-36 profile hidden Markov model spanning 424 match states was built from 83 aligned nematode DAF-36 orthologs, seeded from *C. elegans* DAF-36 (C12D8.5) (22), and searched against every proteome with pyhmmer 0.12.1 (23), a Python binding to HMMER3 (24). Candidate orthologs were then validated by reciprocal best hit against the complete *C. elegans* proteome using BLASTP from BLAST+ 2.12.0 (25). The two criteria were combined explicitly. A species was scored DAF-36-present only when the profile search returned a significant hit and the best reciprocal hit of that protein in *C. elegans* was C12D8.5; absent when neither criterion was met; and unresolved when the two criteria disagreed. Proteome completeness was quantified independently of the DAF-36 result. All 120 proteomes were assessed with BUSCO v6.1.0 (26) in protein mode against the 3131 markers of the nematoda_odb10 lineage set (27), run offline against a local copy of the dataset. Completeness is reported as BUSCO’s complete fraction (single copy plus duplicated). The *C. elegans* proteome scored 99.97% complete (3130 of 3131 markers), validating the procedure. Comparisons of continuous variables between trophic or phylogenetic groups were performed using two-sided Mann–Whitney U tests. Because these comparisons were exploratory, no adjustment was made for multiple testing. Statistical analyses were performed in Python 3.11; sequence manipulation and related analyses used Biopython 1.87.

### *H. glycines* egg and J2 isolation, lipid extraction and HPLC fractionation

The soybean cyst nematode (*H. glycines* Ichinohe) PA3, HG type 0 (race 3) inbred population originally developed by Prakash Arelli from a field population collected from Ames Plantation near Grand Junction, TN by Lawrence Young in 1987 (28), was maintained in the greenhouse at the University of Georgia on the susceptible soybean cultivar Williams 82 and turned over monthly. To obtain eggs, infested roots from ∼30-day-old pot cultures were soaked in water to remove the soil without dislodging the SCN cysts. The roots were placed onto a no. 20-mesh sieve nested on top of a no. 60-mesh sieve. Using a high-pressure hand water sprayer, the cysts were dislodged and collected on the no. 60-mesh sieve. A drill press fitted with a rubber stopper (29) was used to grind the cysts on the no. 60-mesh sieve over a no. 200-mesh sieve nested on a no. 500-mesh sieve. Eggs collected from the no. 500-mesh sieve were further purified by sucrose centrifugal flotation (30), followed by washing several times with water before freezing at -80 °C. To collect pre-parasitic J2s, the eggs were surface-sterilized in 2% sodium azide, rinsed several times with water, and set up to hatch on water containing gentamicin sulfate (22.5 mg/ml) and nystatin (1.5 mg/ml) at 27 °C. After 48 h of hatching, juveniles were collected in siliconized 1.5 mL microfuge tubes by low-speed centrifugation (8,000 rpm) in a tabletop centrifuge and washed several times with water before freezing at -80 °C.

Eggs and J2s were washed extensively with sterile water before lipid extraction. For each preparation, 8 mL of egg pellet or 0.5 mL of J2 pellet was used. Eggs were suspended in 0.9% (w/v) NaCl and homogenized on ice at 20,000 rpm for 15 × 1-min cycles using a Fisherbrand Homogenizer 850, whereas J2s were suspended in 0.9% (w/v) NaCl and sonicated on ice at 60% amplitude for 15 × 10-s pulses. Total lipids were then extracted using a modified Folch method (chloroform:methanol, 2:1, v/v) as previously described (8). Following phase separation, the organic fraction was recovered, dried under reduced pressure, and purified by silica solid-phase extraction (Sep-Pak Silica, Waters Corp.). Lipid extracts were fractionated by preparative reverse-phase HPLC using a Kinetex EVO C_18_ column (250 × 21.2 mm, 5 μm; Phenomenex) on a Shimadzu LC-20AR HPLC system (Shimadzu). Water and acetonitrile containing 0.1% (v/v) formic acid were used as mobile phases at a flow rate of 7.9 mL/min, with acetonitrile increasing from 50% to 100% over 30 min followed by continued elution to 72 min. Fractions were collected at 1-min intervals, dried, resuspended in ethanol, and screened for DAF-12 agonist activity using the cell-based reporter assay.

### Untargeted UHPLC-HRMS/MS analysis

Active fractions from *H. glycines* were separated chromatographically using a Thermo Vanquish UHPLC system using a linear gradient of water (mobile phase A) and acetonitrile (mobile phase B), each containing 0.1% formic acid. Separation was achieved by running a linear ramp of solvent B over a Waters HSST3 C_18_ column (150 mm x 2.1 mm, 1.8 μM). The gradient used for analysis was as follows: 0 – 15 minutes, 80% to 100% mobile phase B; 15 – 17 minutes, 100% mobile phase B flowed isocratically; 17 – 17.1 minutes, 100% to 80% mobile phase B; 17.1 – 23 minutes, 80% mobile phase B flowed isocratically. The flow rate for the gradient was held steady over the course of the run at 0.25 mL/min.

HRMS/MS data were acquired using a data-dependent acquisition method running on a Thermo Scientific Q Exactive HF-X mass spectrometer as reported previously (31). After collection, data were analyzed using Compound Discoverer 3.5 focusing specifically on retention time windows corresponding to subfractions where activity was observed in the reporter assay. Untargeted analysis was performed on active subfractions and adjacent inactive subfractions and all peaks that were significantly increased in active fractions compared to adjacent inactive fractions were recorded. This form of unbiased analysis was performed on a total of four independent preparations and *m/z* 441.2031 was the only peak identified in repeated independent analyses.

### Immunoblotting

COS-7 cells were transiently transfected with pCMX plasmids encoding GR-DAF-12 LBD chimeras using X-tremeGENE HP DNA Transfection Reagent (Roche). Twenty-four hours after transfection, the medium was replaced and cells were cultured overnight before lysis in RIPA buffer (20 mmol/L Tris-HCl, pH 8.0, 150 mmol/L NaCl, 1% (v/v) NP-40, 0.5% (w/v) sodium deoxycholate, 0.1% (w/v) SDS, 20 U/mL DNase I, and 100 µg/mL RNase A) containing protease inhibitors (#11873580001, Roche). Clarified lysates were separated by SDS-PAGE and transferred to nitrocellulose membranes. GR-DAF-12 chimeras were detected with a monoclonal antibody against the glucocorticoid receptor (#66904-1-Ig, Proteintech Group Inc, 1:2000 dilution), with β-actin (#60008-1-Ig, Proteintech Group Inc, 1:2000 dilution) serving as a loading control. Immunoreactive proteins were detected using HRP-conjugated secondary antibodies (#1721011, Bio-Rad) and enhanced chemiluminescence (#34095, Thermo Fisher Scientific) and imaged using a ChemiDoc MP Imaging System (#12003154, Bio-Rad).

## Results

### Plant-parasitic nematode DAF-12 orthologs display differential responsiveness to Δ7-DA

Previous studies established that DAs activate DAF-12 from multiple animal-parasitic nematodes (3). To determine whether DA responsiveness extends to plant-parasitic nematodes, we analyzed DAF-12 orthologs from six representative species spanning diverse parasitic lifestyles within nematode Clade 12 (32). These included three sedentary endoparasites: the cyst nematodes *H. glycines* and *Globodera rostochiensis*, which infect soybean and potato, respectively, and the root-knot nematode *Meloidogyne incognita*, a broad-host-range parasite of numerous crop plants. We also examined three migratory endoparasites: *Ditylenchus destructor*, which infects underground storage organs, including tubers and bulbs, *Aphelenchoides besseyi*, a foliar parasite of rice and other grasses, and *Bursaphelenchus xylophilus*, which infects pine trees. These species were selected to represent diverse host associations among plant-parasitic nematodes.

The LBDs of these receptors along with that of *C. elegans* as a positive control were fused to the amino terminus and DNA-binding domain (DBD) of the GR and tested for Δ7-DA responsiveness in a cell-based reporter assay. All the chimeric receptors were expressed at comparable levels in transfected COS-7 cells (Fig. 1A) and, as expected, Δ7-DA was a potent activator of the *C. elegans* DAF-12 chimera (Fig. 1B). Among the plant-parasitic nematodes, Δ7-DA activated the LBDs of DAF-12 orthologs from *B. xylophilus*, G*. rostochiensis*, *H. glycines* and *A. besseyi* but had little activity on the LBDs from *M. incognita* and *D. destructor* (Fig. 1B). Δ7-DA was most potent and efficacious on the DAF-12 LBD from *B. xylophilus* (EC_50_ ∼100 nM) and showed similar potency on the LBDs from *G. rostochiensis*, *A. besseyi*, and *H. glycines* (EC_50_ values of 0.3–1 μM) (Fig. 1B). Together, these data suggest that while Δ7-DA may be the endogenous ligand for DAF-12 in some plant-parasitic nematode species, it is unlikely to serve this function in others.

**Figure 1.**
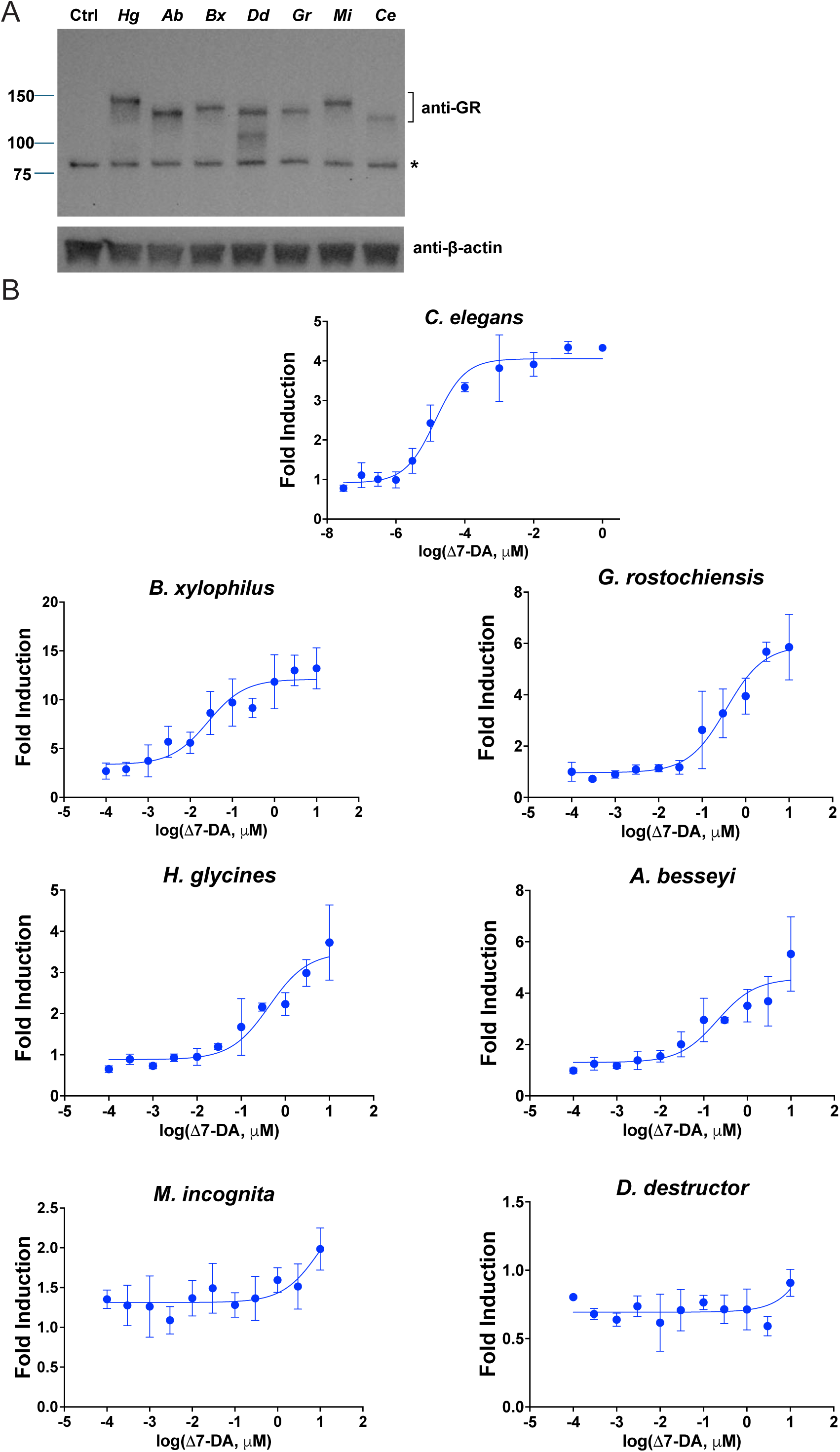
Δ7-Dafachronic acid activates a subset of plant-parasitic nematode DAF-12 orthologs. **(A)** Expression in transiently transfected COS-7 cells of chimeric receptors in which the DAF-12 ligand-binding domain (LBD) of *H. glycines* (Hg), *A. besseyi* (Ab), *B. xylophilus* (Bx), *D. destructor* (Dd), *G. rostochiensis* (Gr), *M. incognita* (Mi) or *C. elegans* (Ce) was fused to the amino terminus and DBD of the glucocorticoid receptor (GR). Blots were probed with antibodies against the GR or β-actin as a loading control. Mock-transfected COS-7 cells lacking plasmid DNA served as the control (Ctrl). The bracket indicates the GR-DAF-12 LBD chimeras. Molecular weight markers (kDa) are shown on the left. The predicted molecular weights of the chimeric proteins are 135 kDa (*Hg*), 127 kDa (*Ab*), 131 kDa (*Bx*), 128 kDa (*Dd*), 131 kDa (*Gr*), 138 kDa (*Mi*) and 125 kDa (*Ce*). A non-specific band is indicated by an asterisk. **(B)** Δ7-DA dose-response curves for the GR-DAF-12 LBD chimeras of the indicated species. Data are fold induction relative to vehicle and are mean ± SD.

### Sequence and structural features associated with DA responsiveness

Comparison of plant-parasitic nematode DAF-12 LBD sequences with structurally characterized DA-responsive receptors revealed conservation of several key ligand-contacting residues (Fig. 2A). All six orthologs retained the arginine corresponding to R599 of *S. stercoralis* DAF-12, which anchors the Δ7-DA C27 carboxylate (9). In contrast, the glutamine that recognizes the C3-keto group (Q637 in *S. stercoralis*) was replaced by serine in *A. besseyi* (S784), despite this receptor retaining Δ7-DA responsiveness. Conversely, the essentially nonresponsive *M. incognita* and *D. destructor* receptors retained both the canonical arginine and glutamine. Thus, conservation of individual ligand-contacting residues alone does not predict Δ7-DA responsiveness.

**Figure 2.**
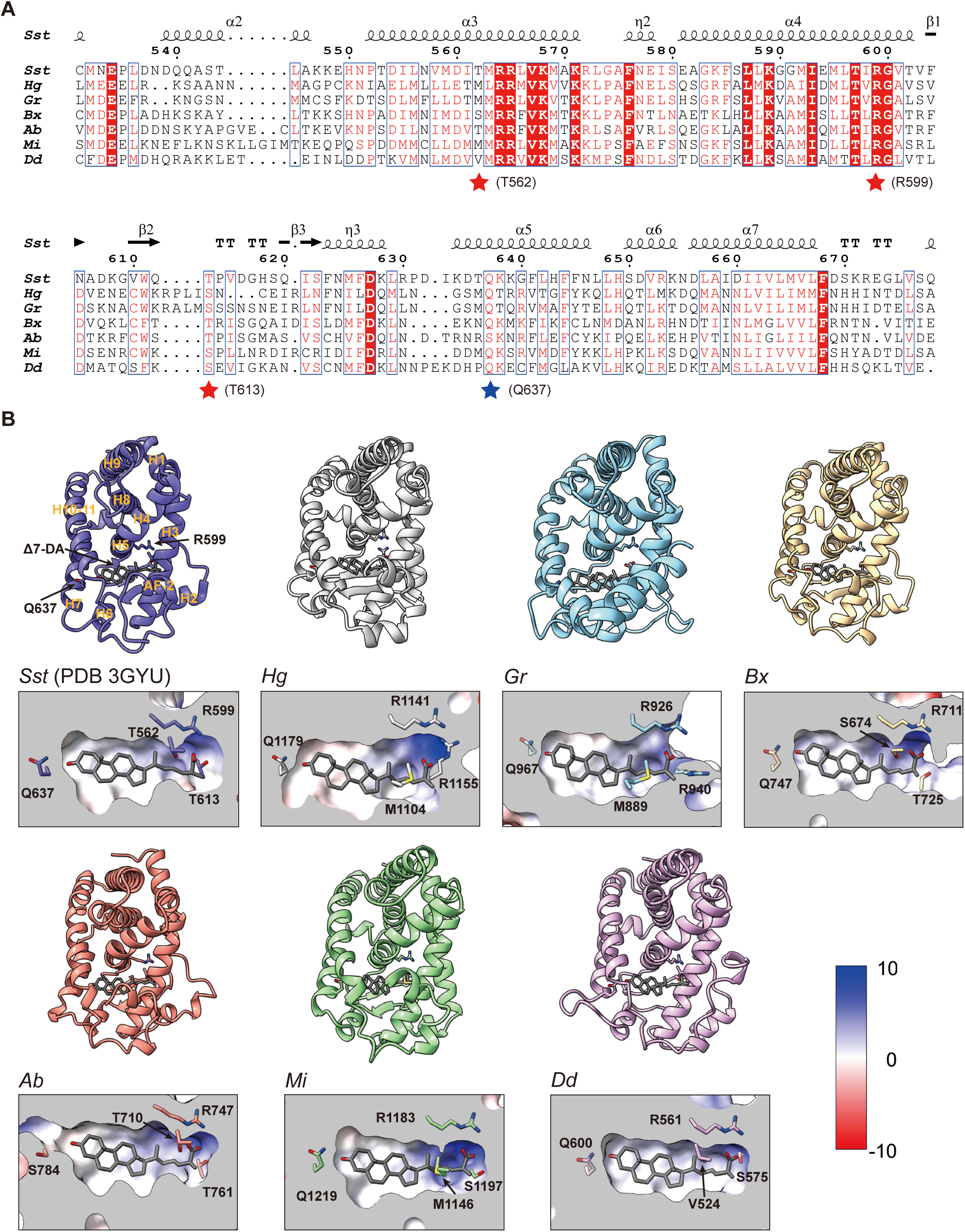
Structural comparison of DAF-12 ligand-binding pockets reveals a combinatorial recognition network underlying differential Δ7-DA responsiveness. **(A)** Structure-guided sequence alignment of the ligand-binding domains (LBDs) of DAF-12 from *S. stercoralis* (*Sst*; PDB 3GYU), *H. glycines* (*Hg*), *G. rostochiensis* (*Gr*), *B. xylophilus* (*Bx*), *A. besseyi* (*Ab*), *M. incognita* (*Mi*) and *D. destructor* (*Dd*). Amino acid numbering and secondary-structure elements are based on the *Sst* DAF-12 structure, with α- and 3_10_-helices (η) shown as ribbons, β-strands as arrows, and turns indicated by T. Conserved residues are highlighted in red. Red stars indicate residues corresponding to the C27-carboxylate recognition network and the blue star marks the residue adjacent to the C3 keto-recognition site. The overall architecture of the ligand-binding pocket is highly conserved, whereas substitutions at key ligand-contacting positions define a combinatorial recognition network associated with differential Δ7-DA responsiveness. **(B)** Structural comparison of the DAF-12 LBDs from *Sst*, *Hg*, *Gr*, *Bx*, *Ab*, *Mi* and *Dd*. The LBDs were predicted using AlphaFold3, and Δ7-dafachronic acid (Δ7-DA) was redocked into each model. Δ7-DA is shown as gray sticks. Electrostatic surface representations highlight residues surrounding the ligand-binding pocket. In the active *Sst* reference structure, the C27 carboxylate is coordinated by T562, R599 and T613, whereas the C3 keto group is recognized by Q637. Although Δ7-DA adopts a broadly similar pose in all models, variations in the carboxylate-recognition network distinguish the responsive receptors *Hg*, *Gr*, *Bx* and *Ab* from the nonresponsive receptors *Mi* and *Dd*.

To identify structural features that might explain these differences, we generated AlphaFold3 models of all six plant-parasitic DAF-12 LBDs and docked Δ7-DA into each model using the Δ7-DA-bound *S. stercoralis* DAF-12 LBD (PDB 3GYU) as the active-state reference (Fig. 2B). All six models adopted the canonical nuclear receptor fold and accommodated Δ7-DA in a broadly similar orientation without major steric clashes, indicating that differential responsiveness is unlikely to result from gross differences in pocket architecture. Instead, structure-guided alignment revealed systematic variation in the network of residues surrounding the ligand carboxylate (Fig. 2A, 2B).

In the *S. stercoralis* structure, the Δ7-DA C27 carboxylate is coordinated by T562, R599, and T613 (9). The more strongly responsive *B. xylophilus* and *A. besseyi* receptors retain similar S/T–R–T networks (S674–R711–T725 and T710–R747–T761, respectively). *H. glycines* and *G. rostochiensis* instead contain M–R–R configurations (M1109–R1146–R1160 and M889–R926– R940), introducing a second arginine that could provide an alternative multipoint interaction with an acidic ligand. By contrast, the nonresponsive *M. incognita* and *D. destructor* receptors contain M/V–R–S configurations (M1146–R1183–S1197 and V524–R561–S575), in which substitution of threonine by the shorter serine side chain may weaken carboxylate coordination (Fig. 2A, 2B). Together with the lack of correlation between the C3-keto-contacting glutamine and receptor activity, these observations suggest that Δ7-DA responsiveness is a combinatorial property of the ligand-binding pocket rather than a consequence of any single ligand-receptor interaction.

These structural differences may influence not only ligand recognition but also coupling of ligand binding to receptor activation. Nuclear receptor agonism requires stabilization of the C-terminal AF-2/H12 surface that recruits LxxLL-containing coactivators, and alterations in residues surrounding the ligand-binding pocket could impair this coupling even when Δ7-DA can be accommodated. The modest responsiveness of *H. glycines* DAF-12 may represent an intermediate configuration: its primary ligand-contacting residues are retained, but the M1109–R1146– R1160 carboxylate-recognition network differs substantially from that of DA-responsive receptors and may instead favor an endogenous acidic ligand distinct from Δ7-DA (see below). Collectively, these findings indicate that DA responsiveness depends on the broader structural network that coordinates ligand recognition with formation of the active AF-2 surface.

### Evidence that DA is not the endogenous ligand for *H. glycines* DAF-12

Given the differential responsiveness of plant-parasitic nematode DAF-12 receptors to Δ7-DA, we examined their evolutionary relationships with DAF-12 proteins from other nematodes. Maximum-likelihood phylogenetic analysis revealed that DAF-12 LBDs from plant-parasitic nematodes clustered predominantly together, consistent with their placement in Clade 12 (Fig. 3A, S2) (32). Interestingly, however, analysis of enzymes involved in DA biosynthesis revealed that DAF-36, which catalyzes the first committed step in Δ7-DA biosynthesis from cholesterol (22), was absent from nearly all Clade 12 nematodes examined, including 16 of 17 plant-parasitic nematode species (Fig. 3B). This lineage-associated pattern was supported by the complete phylogeny and was unlikely to reflect differences in genome completeness (Figs. S2 and S3). The widespread absence of DAF-36 in plant-parasitic nematodes suggests that Δ7-DA is unlikely to serve as their endogenous DAF-12 ligand.

**Figure 3.**
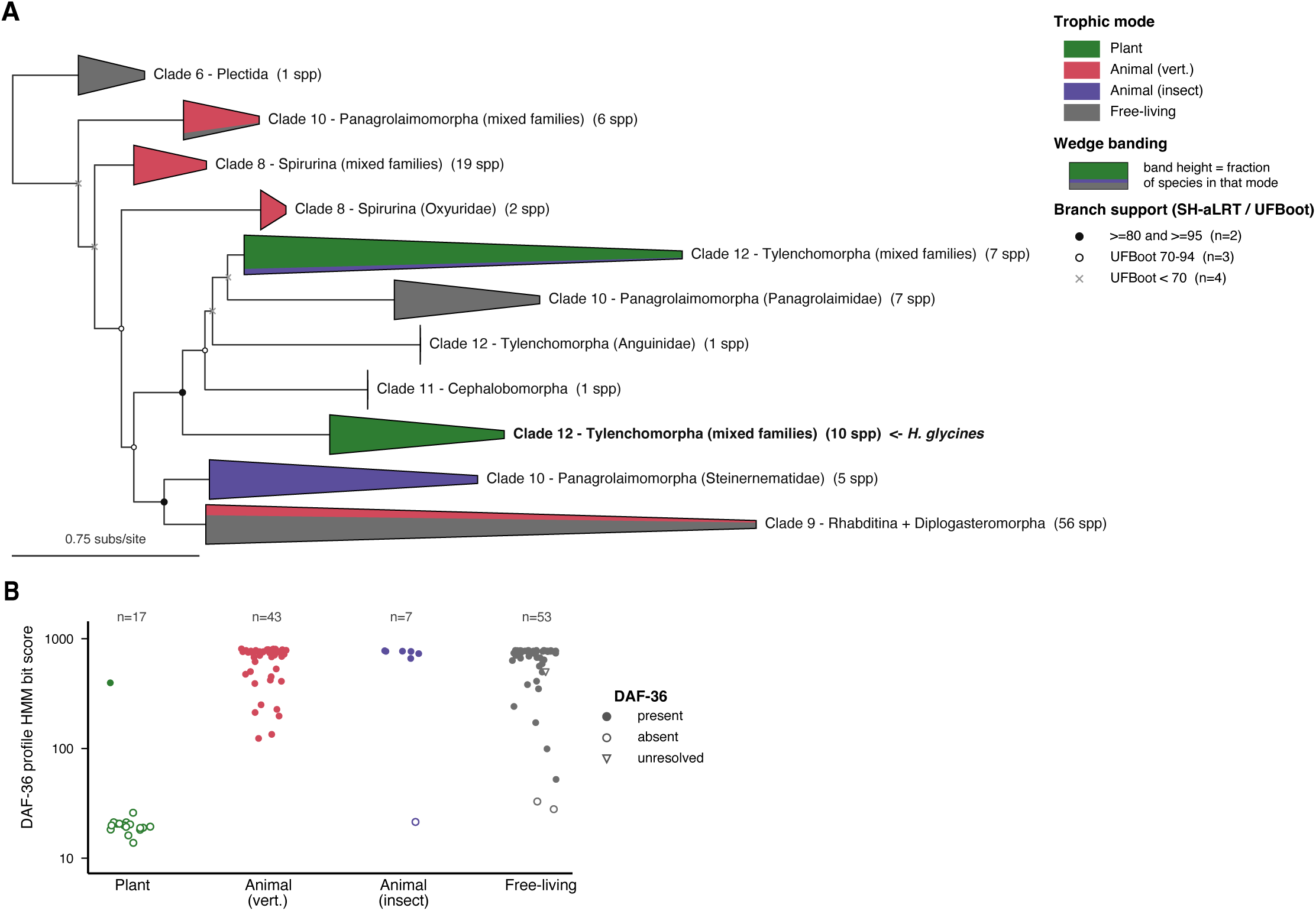
DAF-12 ligand-binding domains and dafachronic acid synthesis enzyme DAF-36 suggest ligand diversification in plant-parasitic nematodes. **(A)** Simplified maximum-likelihood tree based upon phylogenetic analysis of DAF-12 ligand-binding domains across nematodes. Major nematode lineages are indicated by shaded blocks and labeled according to nematode taxonomy, with the corresponding molecular clades shown where applicable. Plectina serves as the outgroup; Spirinuria contains representatives of Clade 8; Rhabditina contains Clade 9 nematodes, including *C. elegans* and many animal parasites; and Tylenchina contains diverse Clade 10, 11 and 12 nematodes. Within Tylenchina, Tylenchoidea includes the agriculturally important cyst (*Heterodera* and *Globodera*) and root-knot (*Meloidogyne*) nematodes. Node support information is indicated in the figure. **(B)** Most plant-parasitic nematodes appear to lack DAF-36 homologs. Y-axis indicates bit score from profile HMM search against *C. elegans* DAF-36. The best match from the profile HMM search was verified by reciprocal BLAST.*n* indicates the number of species per trophic group.

Since *H. glycines* is among the most devastating plant-parasitic nematodes, we searched for its endogenous DAF-12 ligand. *H. glycines* develops through four juvenile stages before reaching adulthood (33). The first-stage juvenile molts within the egg and the J2 emerges as the infective stage, migrates through the soil and penetrates soybean roots. Thus, eggs and J2s span the developmental transitions likely to be regulated by DAF-12. Lipids extracted from *H. glycines* eggs or J2s were fractionated by preparative HPLC, and individual fractions were tested for activity on the LBDs of *H. glycines* and the five other plant-parasitic DAF-12 orthologs.

Multiple HPLC fractions from egg extracts, including fractions 28, 30, 34, and 36, activated the *H. glycines* DAF-12 LBD, whereas activity in J2 extracts was predominantly detected in fraction 36 (Fig. 4A, 4B). Importantly, none of the three known endogenous DAs, namely Δ4-DA, Δ7-DA and Δ1,7-DA, was detected in either egg or J2 extracts by high-resolution mass spectrometry. DA activity was also not detected in egg fractions using the sensitive *C. elegans* DAF-12 LBD cell-based reporter assay (Fig. 4C). Interestingly, the same egg-derived fractions that activated *H. glycines* DAF-12 also activated *G. rostochiensis* DAF-12, but not DAF-12 from the other plant-parasitic nematodes (Fig. 4A), suggesting that these two confamilial cyst nematodes may share common endogenous ligands. *H. glycines* and *G. rostochiensis* are also the only two orthologs in this set that carry the M–R–R carboxylate-recognition network (Fig. 2A), providing a structural rationale for their shared and selective response to the egg-derived fractions.

**Figure 4.**
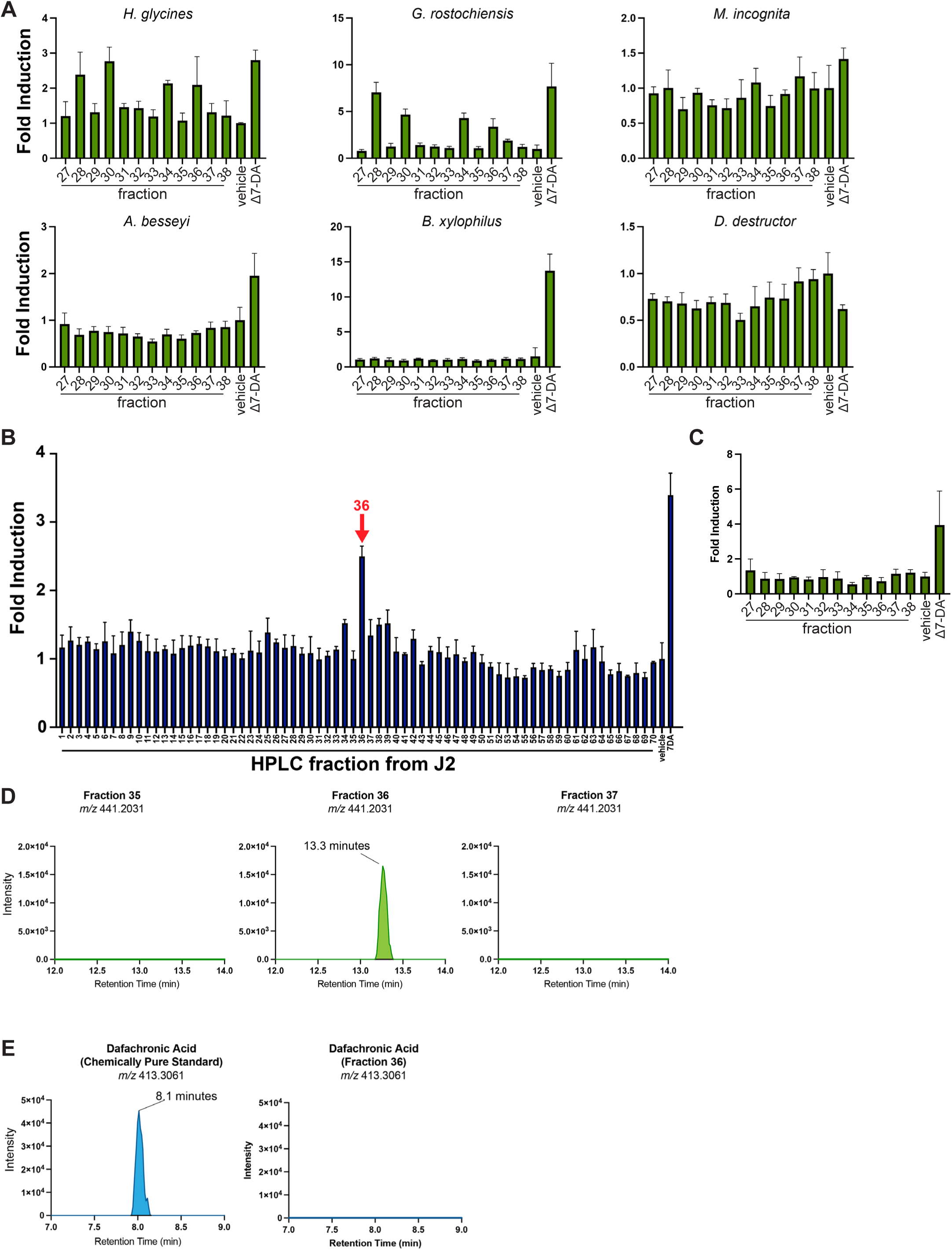
*H. glycines* egg and J2 extracts have DAF-12 activity that is distinct from dafachronic acid. **(A)** Lipid fractions from *H. glycines* eggs were assayed for transcriptional activity on each of the six plant-parasitic GR-DAF-12 chimeric receptors. **(B)** Lipid fractions from *H. glycines* J2s were assayed on the *H. glycines* GR-DAF-12 LBD chimera. Ethanol vehicle and Δ7-DA (1 μM) were included as controls. **(C)** Lipid fractions from *H. glycines* eggs were assayed on the *C. elegans* DAF-12 LBD chimera. Ethanol vehicle and Δ7-DA (1 μM) were included as controls. All cell-based transfection data are fold induction relative to vehicle and are mean ± SD. **(D)** Untargeted UHPLC-HRMS/MS analysis of fractions 35-37 from *H. glycines* extracts reveals a unique molecular feature in fraction 36. Using electrospray ionization, we identified *m/z* 441.2031 with a retention time of 13.3 minutes. **(E)** Analysis of Δ7-DA using electrospray ionization with the same analytical method demonstrates that it elutes at a retention time distinct from *m/z* 441.2031 and is not present in fraction 36.

Because fraction 36 from both egg and J2 extracts had *H. glycines* DAF-12 activity, it was selected for further analysis. Orthogonal UHPLC fractionation localized activity to a single subfraction, defining the retention time window for untargeted LC-MS/MS analysis. Comparison of active and adjacent inactive fractions identified a single ion (*m/z* 441.2031, negative mode) that reproducibly co-eluted with DAF-12 activity and was enriched 5- to 7-fold in active fractions (Fig. 4D). This ion eluted approximately six minutes later than authentic Δ7-DA, which was not detected in fraction 36 (Fig. 4E), further indicating that the active species is distinct from DA. The ion was consistent with loss of a proton (M-H), suggesting the presence of an acidic functional group analogous to the DA carboxylate. Data supporting additional adduct formation were not observed in our spectra. Although definitive structural identification will require substantial scale-up, these findings indicate that *H. glycines* produces an endogenous DAF-12 agonist chemically distinct from the known DAs.

## Discussion

The discovery of the DAF-12 receptor and DA as its endogenous ligand transformed our understanding of *C. elegans* and animal-parasitic nematode development by establishing that their life-cycle progression is regulated by an endocrine signaling pathway analogous to steroid hormone signaling in vertebrates (3). The current study extends these discoveries to plant-parasitic nematodes and reveals that the evolutionary history of DAF-12 signaling is more complex than previously appreciated. We demonstrate that DAF-12 orthologs from diverse plant-parasitic nematode species exhibit markedly different responses to Δ7-DA. Whereas receptors from *B. xylophilus*, *G. rostochiensis*, *H. glycines*, and *A. besseyi* retained responsiveness, receptors from *M. incognita* and *D. destructor* were largely unresponsive despite conserving the overall architecture of the LBD. Comparative structural modeling suggests that relatively modest structural changes within the ligand-binding pocket are sufficient to alter ligand responsiveness while preserving receptor function. Rather than a single determinant, responsiveness tracks with a combinatorial network of residues that surrounds the ligand carboxylate and couples it to the receptor’s coactivator-binding surface.

The most significant finding of this work is that the endogenous ligand for *H. glycines* DAF-12 is highly unlikely to be DA. Egg and J2 extracts lacked detectable DA yet contained lipophilic fractions that activated *H. glycines* DAF-12. Bioassay-guided fractionation coupled with untargeted LC-MS/MS further identified a candidate acidic molecule that co-eluted with receptor activity but exhibited chromatographic properties distinct from DA. Although the structure of this molecule remains unknown, these findings strongly suggest that *H. glycines* synthesizes an endogenous DAF-12 agonist that is chemically distinct from the known DAs. The observation that the same *H. glycines* egg fractions also activate *G. rostochiensis* DAF-12 raises the intriguing possibility that these cyst nematodes share a new class of DAF-12 ligands. Our models provide a structural correlate for this shared pharmacology in the M–R–R carboxylate-recognition network unique to these two receptors and predict that the endogenous ligand carries an acidic group positioned to engage the additional arginine.

Collectively, our findings support a model in which DAF-12 has been evolutionarily conserved whereas its endogenous ligands have diversified. This conclusion is further supported by the apparent absence of DAF-36, which catalyzes the first committed step in Δ7-DA and Δ1,7-DA biosynthesis from cholesterol, in most plant-parasitic nematodes. Adaptation to different hosts and environments may have favored the evolution of novel DAF-12 ligands in these species while preserving the receptor as a central developmental switch. Because plant-parasitic nematodes acquire sterols primarily from phytosterols such as β-sitosterol (34), their DAF-12 ligands may be derived from plant sterol precursors rather than cholesterol. In *H. glycines*, whether ligand biosynthesis involves C24 dealkylation of phytosterols, as reported for *C. elegans* and several plant-parasitic nematodes (34), or instead retains the phytosterol side chain remains to be determined. The continued responsiveness of several plant-parasitic DAF-12 orthologs to Δ7-DA suggests that the transition from DA to alternative ligands has occurred to varying degrees across nematode lineages rather than through a single evolutionary event.

These findings could have important translational implications. Consistent with the role of DAF-12 in regulating developmental progression in animal-parasitic nematodes, pharmacologic activation of DAF-12 in *S. stercoralis* suppresses formation of infective larvae *in vitro* and reduces parasite burden in infected gerbils (8), providing proof of principle that this pathway is therapeutically tractable. Plant-parasitic nematodes cause billions of dollars in agricultural losses annually and represent a growing threat to global food security, yet relatively few selective nematicides are available. Manipulating DAF-12 signaling with agonists or antagonists is therefore an attractive strategy for controlling these pathogens. Moreover, the apparent divergence of endogenous DAF-12 ligands among plant-parasitic nematodes raises the possibility of developing highly selective control strategies tailored to individual species. The structural models presented here provide a starting point for such efforts by mapping the pocket positions that differ among orthologs and by identifying coupling to the AF-2 surface as a second, largely unexplored point of intervention.

Finally, this work is dedicated to the memory of David Mangelsdorf, who initiated this project and whose vision established the conceptual framework on which it is built. The discovery of DAs and the demonstration that DAF-12 signaling regulates developmental transitions across nematode species fundamentally changed the field. By extending these discoveries to plant-parasitic nematodes, we show that while DAF-12 is conserved, the endogenous hormones that regulate its activity have likely diversified in response to the distinct ecological pressures encountered by the different species. Defining these novel ligands will not only deepen our understanding of nematode biology but also provide new opportunities for protecting crops from some of the world’s most destructive pathogens.

## Declaration of Interest, Funding and Acknowledgements

None of the authors declare conflicts of interest that could be perceived as prejudicing the impartiality of the research reported. This work was supported by the NIH (R01AI167967 to J.J.C.), the Robert A. Welch Foundation (I-1948-20240404 to J.J.C., I-1275 to D.J.M.), Lyda Hill Philanthropies (Hill Prize to S.A.K. and D.J.M.) and the Howard Hughes Medical Institute (J.J.C. and D.J.M.). L.G.Z., T.P.M. and the Children’s Research Institute Metabolomics Facility are supported by an award from the Cancer Prevention Research Institute of Texas (CPRIT Core Facilities Support Award RP24094).

Artificial intelligence tools were used in both data analysis and manuscript preparation. Claude Science (beta; Opus 5 with Sonnet 5 as reviewer) was used to assemble sequences, run tools IQ-TREE 2, BUSCO 6.1.0, pyhmmer 0.12.1, and BLAST+ 2.12.0, and generate Figures 3, S2, and S3. Following the initial production of IQ-TREE, BUSCO, and pyhmmer outputs, the model was asked to generate a protocol, delete all of the outputs, and repeat the analysis using the protocol it generated. The maximum-likelihood tree shown in Figures 3A and S2 was validated manually with IQ-TREE 2 and the absence of DAF-36 homologs was validated manually with BLASTP. ChatGPT (GPT-5.6 Sol; OpenAI) was used for language editing. All AI-generated analyses and text were reviewed by the authors, who take full responsibility for the accuracy and content of the publication.

---

**Supplementary Figure S1.**
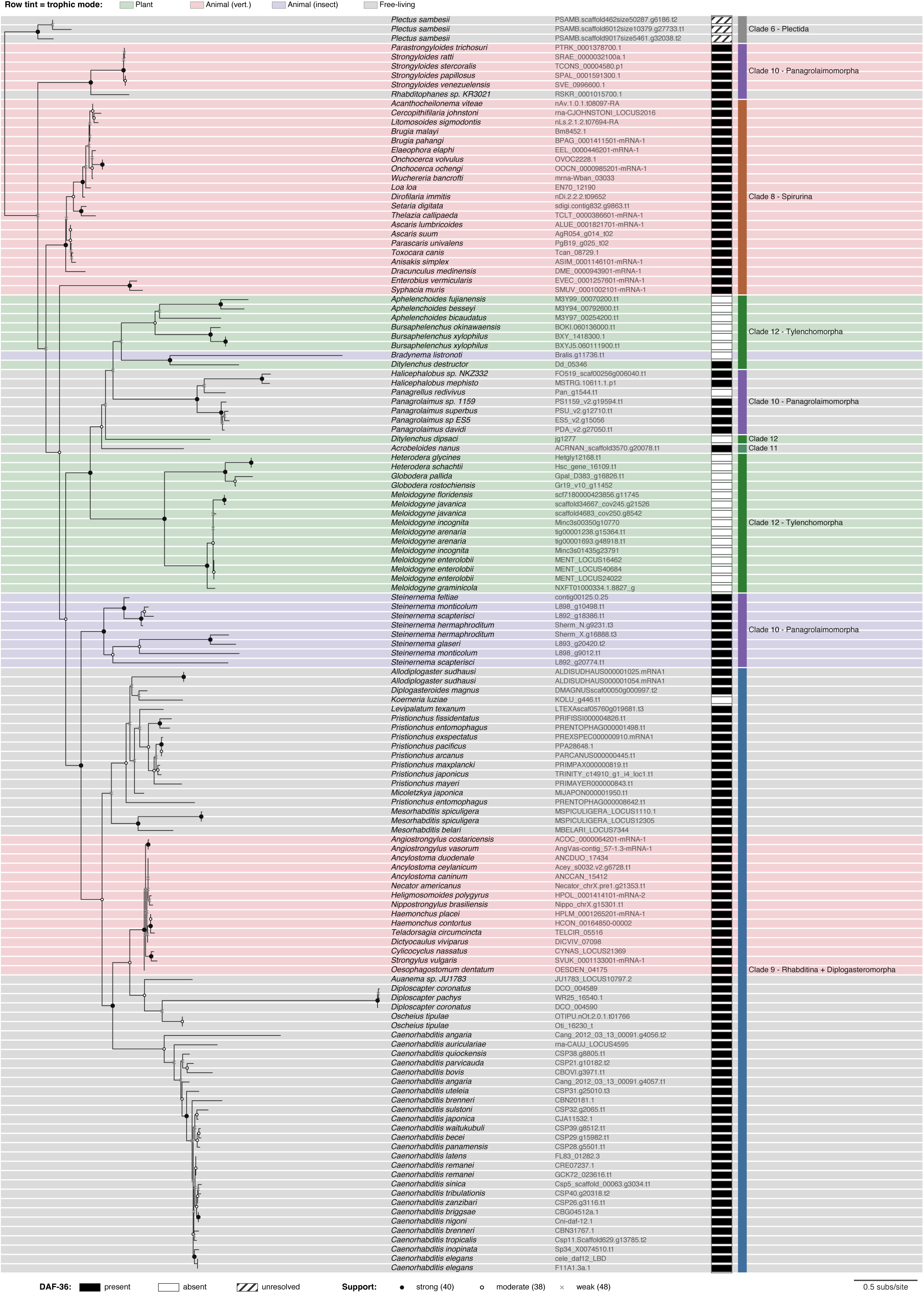
Hinge and LBD amino acid sequences of plant-parasitic nematodes that were incorporated into the chimeras.

**Supplementary Figure S2.**
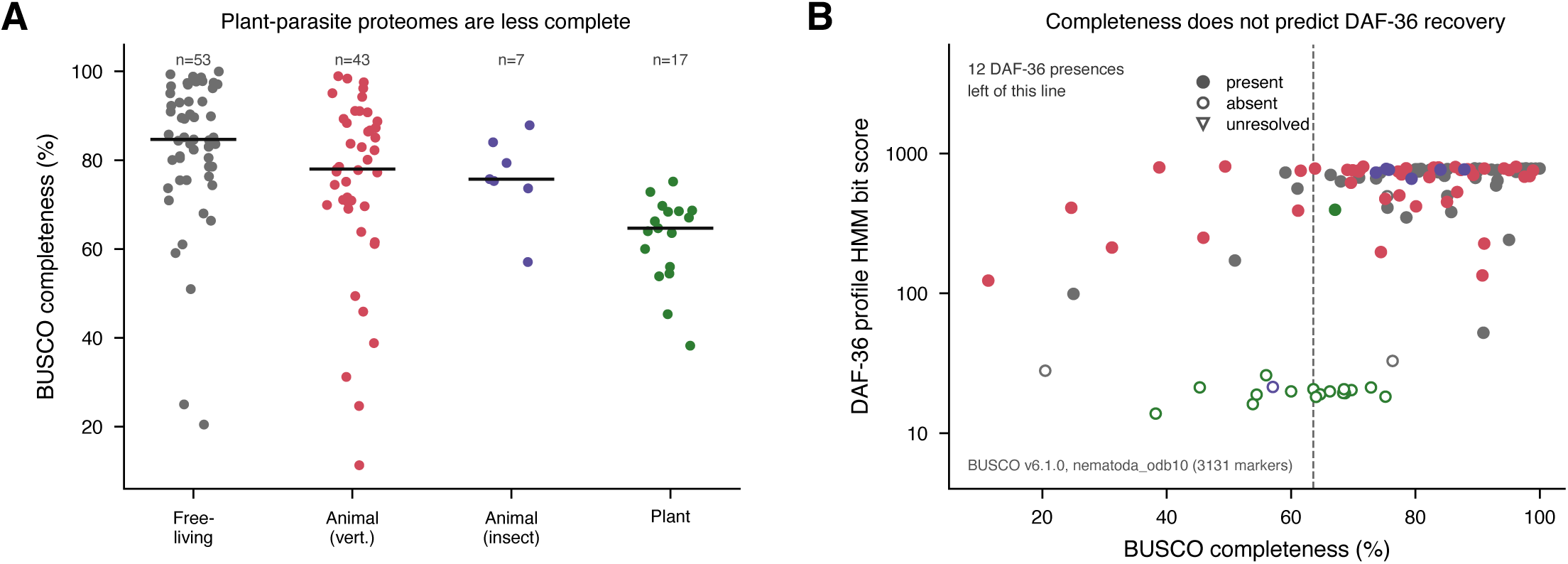
Complete maximum-likelihood tree from phylogenetic analysis of DAF-12 ligand-binding domain across 115 species of nematodes. Node support information is indicated in the figure. The right side of the graph indicates whether a DAF-36 homolog was detected as determined by a reciprocal BLAST search. Newick tree is provided in File S1.

**Supplementary Figure S3. Genome completeness does not explain lack of DAF-36 homologs in plant-parasitic nematodes. (A)** Graph indicating BUSCO completeness of genomes from the nematodes examined split by their trophism. **(B)** DAF-36 profile HMM bit score plotted against BUSCO completeness. Several nematodes with detectable DAF-36 homologs have lower BUSCO scores than the plant parasitic nematodes. Blue empty circle near the bottom indicates *H. glycines*.

